# *Unraveling the Potential Role of Tecomella undulata* in Experimental NASH

**DOI:** 10.1101/2022.08.26.505375

**Authors:** Akshatha N. Srinivas, Diwakar Suresh, Deepak Suvarna, Pankaj Pathak, Suresh Giri, Suman, Suchitha Satish, Saravana Babu Chidambaram, Divya P. Kumar

**Affiliations:** Department of Biochemistry, CEMR, JSS Medical College, JSS Academy of Higher Education and Research, Mysuru, Karnataka, India; Department of Gastroenterology, JSS Medical College and Hospital, JSS Academy of Higher Education and Research, Mysuru, India; JSS Ayurveda Medical College and Hospital, Mysuru, Karnataka, India; Zydus Research Centre, Cadila Healthcare Limited, Ahmedabad, Gujarat, India; Government Ayurveda Medical College, Mysuru, Karnataka, India; Department of Pathology, JSS Medical College and Hospital, JSS Academy of Higher Education and Research, Mysuru, India; Department of Pharmacology, JSS College of Pharmacy, JSS Academy of Higher Education and Research, Mysuru, India

**Keywords:** Insulin resistance, hepatic inflammation, oxidative stress, ER stress, herbal medicine

## Abstract

**Background and Aim:** The pathophysiology of NASH is complex owing to its diverse pathological drivers, and until recently, there were no approved drugs for this disease. *Tecomella undulata* is a popular herbal medicine used to treat hepatosplenomegaly, hepatitis, and obesity. However, the potential role of *Tecomella undulata* in NASH has not yet been scientifically investigated.

**Experimental Procedure:** Mice fed with chow diet and normal water (CDNW) or western diet and sugar water (WDSW) for 12 weeks were randomized to receive vehicle control, Saroglitazar, or *Tecomella undulata* for an additional 12 weeks. Insulin resistance, lipid profiles, histological analysis, and liver enzymes were assessed. The oxidative stress, ER and inflammatory markers were determined by quantitative RT-PCR (qRT-PCR) and western blot analysis.

**Results and Conclusion:** The administration of *Tecomella undulata* via oral gavage lowered body weight, insulin resistance, alanine transaminase (ALT), aspartate transaminase (AST), triglycerides, and total cholesterol in WDSW mice but had no effect on CDNW mice. *Tecomella undulata* improved steatosis, lobular inflammation, and hepatocyte ballooning and resolved NASH in WDSW mice. Furthermore, *Tecomella undulata* also alleviated the WDSW-induced ER stress and oxidative stress, enhanced antioxidant status, and thus reduced inflammation in the treated mice. Of note, these effects were on par with Saroglitazar, the approved drug used to treat human NASH and positive control used in the study.

Thus, our findings indicate the potential of *Tecomella undulata* to ameliorate WDSW-induced steatohepatitis, and these preclinical data provide a strong rationale for assessing *Tecomella undulata* for the treatment of NASH in humans.

## 1. INTRODUCTION

Nonalcoholic fatty liver disease (NAFLD) is a multisystemic disease that encompasses a spectrum of conditions, including benign fatty liver, nonalcoholic steatohepatitis (NASH), fibrosis, cirrhosis, and hepatocellular carcinoma (Powell et al., 2021). NAFLD, a significant health burden caused by the rise in obesity and sedentary lifestyle, is estimated to have a 25% global prevalence and is expected to quadruple by 2030 with inadequate management (Younossi et al., 2017). The fatty liver resulting from overnutrition and insulin resistance promotes proinflammatory settings for NASH (Friedman et al., 2018). The liver is overworked as a result of increased hepatic intake, decreased lipolysis, and de novo lipogenesis, which results in lipotoxicity, an inflammatory response, cell death, and fibrogenic activation (Cotter and Rinella, 2020). Thus, despite a plethora of studies, the pathobiology of NASH is poorly understood due to its diverse disease drivers, leading to heterogeneous therapeutic responses. There are still unmet medical needs and challenges, and until recently, there were no approved medications NAFLD. However, in 2020, Saroglitazar became the first medication for NASH to receive approval and is prescribed to patients in India (Shetty et al., 2015).

The current paradigm of multiple hits involving a number of factors acting in parallel provides a better delineation of NASH development and progression (Buzzetti et al., 2016). Oxidative stress is a major contributor among the multiple hits of NAFLD progression. The imbalance between the production and scavenging of reactive oxygen species (ROS) due to hepatic lipid overload leads to metabolic dysfunction followed by an inflammatory response (Chen et al., 2020). The accumulation of unfolded proteins in the endoplasmic reticulum (ER) lumen as a result of lipotoxicity causes ER stress, which is characterized by elevated levels of ER stress markers and mediates the inflammatory cascades in the progression of NAFLD (Kakazu et al., 2016; Ashraf, Sheikh, 2015). In contrast, antioxidant levels are significantly downregulated in NAFLD patients, reiterating the root cause of disease progression (García-Ruiz and Fernández-Checa, 2018). Furthermore, the disease’s progression is determined by the cellular stress-mediated inflammatory tone caused by immune cell crosstalk (Schuster et al., 2018). As a result, the discovery of drugs with pleiotropic effects is an effective therapeutic strategy for combating the disease.

In recent years, plant-based medicines have gained the limelight, along with plant-based dietary patterns. Whole medicinal plants, which are unpurified extracts, have been traditionally used in many parts of the world as an effective therapy for NAFLD management (Li et al., 2021). The holistic approaches to herbal medicine have benefits since they allow for the identification of pure natural ingredients in these medications. *Tecomella undulata* (Sm.) Seem. (Rohitaka) [The plant name has been verified with World Flora Online (http://www.worldfloraonline.org) and MPNS (https://mpns.science.kew.org)] is one such traditional medicine belonging to the Bignoniaceae family which is widely used in Ayurveda for its pharmacological effects including hepatoprotection, immunomodulation, anti-inflammatory and antimicrobial activity (Kumawat et al., 2012; Jain et al., 2012; Thanawala and Jolly, 1993). *Tecomella undulata*, a native shrub of northeastern and western India, has also shown promise as an anti-obesity formulation (Alvala et al., 2013). The stem bark powder of *Tecomella undulata* is widely used in treating ascites with hepatosplenomegaly and also in hepatitis as it is known to work efficiently as a blood purifier (Saggoo et al., 2017). Of note, the stem bark powder of *Tecomella undulata* is a key ingredient in many herbal remedies for the treatment of inflammatory liver illnesses, including Livo-plus, Liv-52, Herboliv, Livfit, Amylcure, Himoliv, Livosan, SLIM-U capsules and Exol (Dhir and Shekhawat 2012). Experimental studies have shown the hepatoprotective potential of *Tecomella undulata* against paracetamol-, carbon tetrachloride-, and thioacetamide-induced hepatotoxicity (Saxena et al., 2021). Thus, the medicinal value with effective pharmacological effects of *Tecomella undulata* has drawn a lot of interest in the liver field. However, the potential role of *Tecomella undulata* and its molecular mechanisms in NASH has not been investigated.

The overall goal of this study was to investigate the pharmacological effects of *Tecomella undulata* and offer solid justification for the clinical treatment of NASH. We here report the effects of *Tecomella undulata* on the pathogenesis of NASH using a mouse model (detailed in the methods section) that mimics the key elements of human NASH in terms of histological progression and molecular signature (Charlton et al., 2011; Asgharpour et al., 2016; Santhekadur et al., 2018). The specific objectives of the study were to demonstrate the efficacy of *Tecomella undulata* on (i) systemic obesity and insulin resistance, (ii) inflammation, and (iii) key pathogenic signaling pathways in NASH. Taken together, these provide novel data demonstrating scientific evidence for the traditional claim of *Tecomella undulata*’s hepatoprotective action in the context of NASH.

## 2. MATERIALS AND METHODS

### 2.1. Chemicals

Saroglitazar was a generous gift from Zydus Research Center, Cadila Healthcare Limited, Ahmedabad, Gujarat, India. Antibodies to JNK, ERK1/2 were procured from Santa Cruz Biotechnology, Inc. (Santa Cruz, CA); mouse insulin ELISA kit from Krishgen Biosystems; ROS Detection Assay Kit from BioVision (MA, USA); TBARS Assay kit from Cayman Chemicals (MI, USA). The western diet was customized and purchased from Research Diets Inc. (NJ, USA); glucose and fructose were procured from Himedia Laboratories (India).

### 2.2. Plant Material

*Tecomella undulata* crude powder was purchased from Ayur Drugs & Pharmaceuticals. As per the details provided by the company, the barks of *Tecomella Undulata* were collected from the natural habitat of Jaipur, Rajasthan. We confirmed the authentication from the JSS Ayurveda Medical College and Hospital and Government Ayurveda Medical College, Mysuru. The plant material was also authenticated by comparison with the reference specimen (EP 427), preserved at the herbarium of National Bureau of Plant Genetic Resources. Furthermore, the stem bark powder of *Tecomella Undulata* was subjected to pharmacopoeia tests.

### 2.3. Cell culture

HepG2 and Huh7 cells were cultured in DMEM (Himedia Laboratories) containing 1 g/L and 4.5 g/L glucose, respectively, supplemented with 10% fetal bovine serum (Gibco), L-glutamine, sodium pyruvate, nonessential amino acids, and 100 U/ml penicillin–streptomycin incubated at 37^0^C in 5% CO_2_.

### 2.4. In vitro Cytotoxicity assay

The WST-1 assay (Takara) was performed on HepG2 cells and Huh7 cells to assess the cellular cytotoxicity of *Tecomella undulata*. Briefly, 1×10^4^ cells were seeded into a 96-well plate and cultured up to 70% confluency. Cells were treated with *Tecomella undulata* powder dissolved in methanol at different concentrations (0, 10, 20, 50, 100, 250, 500, 750, 1000 µg/ml) for 24 h, 48 h, and 72 h. At the end of the treatment, WST-1 assay mix (1:10 dilution) was added to each well and incubated for 1 h at 37^0^C in 5% CO_2_. The absorbance was measured at 450 nm on a multi-mode plate reader (EnSpire™ Multimode Plate Reader, Perkin Elmer). Percent cell viability was calculated in comparison to the control (no treatment).

### 2.5. Animal model

The 6-7 week-old male C57Bl/6 mice (Charles River Laboratories) used in the study were maintained as described previously (Charlton et al., 2011; Tsuchida et al., 2018). All mice were housed on a 12h light-dark cycle in 21-24^0^C animal facility administered by the Centre for Experimental Pharmacology and Toxicology, JSS AHER. The mice were regularly monitored for body weight, food, and water intake and were euthanized at varied time points following dietary and drug interventions. Blood collection and tissue harvesting were performed according to the approved protocols in accordance with the guidelines and regulations of the Committee for the Purpose of Control and Supervision of Experiments on Animals (CPCSEA), Govt. of India (JSSAHER/CPT/IAEC/060/2021).

### 2.6. Study design and interventions

After a week of acclimatization, the mice were fed with either a chow diet (CD) or a western diet (WD) with high fat, high sucrose, and high cholesterol (WD, 21% fat, 41% sucrose, and 1.25% cholesterol by weight, Research Diet, Inc. USA) along with normal water (NW) or high sugar (SW, 18.9 g/L d-glucose and 23.1 g/L d-fructose) containing drinking water ad libitum for 12 weeks. *Tecomella undulata* and Saroglitazar were dissolved in 0.5% carboxymethylcellulose (CMC) and Tween 80 in a 99.5:0.5 ratio and administered via oral gavage once a day for 12 weeks. Saroglitazar was used as a positive control throughout the study. After 12 weeks, mice were randomly divided into 6 groups. Three groups in chow diet fed mice: CDNW with vehicle control-VC (0.5% CMC and Tween 80 in a 99.5:0.5 ratio), CDNW with Saroglitazar (4 mg/kg/day), and CDNW with *Tecomella undulata* (80 mg/kg/day). Three groups in western diet-fed mice: WDSW with vehicle control-VC (0.5% CMC and Tween 80 in a 99.5:0.5 ratio), WDSW with Saroglitazar (4 mg/kg/day), and WDSW with *Tecomella undulata* (80 mg/kg/day). The mice were constantly monitored for food and water intake, and the body weight was recorded on a weekly basis. At the end of treatment (24 weeks), mice were euthanized; blood was collected and biochemical analysis was performed.

### 2.7. Choice of *Tecomella undulata* dosage

Pharmacopoeia tests were performed to ensure the pharmaceutical quality of *Tecomella undulata* stem bark powder for use in *in vivo* research (**Supplementary Table 1**). The dose of *Tecomella undulata* was based on the body surface area (BSA) conversion factor as recommended by the Food and Drug Administration of the US federal government (https://www.fda.gov/media/72309/download). A dose of 80 mg of *Tecomella undulata*/kg body weight of mice was calculated based on the equation: therapeutic human dose of *Tecomella undulata* (6g/kg)× body surface area (BSA) conversion factor for mice (0.0026) × inter species conversion factor (5).

### 2.8. Blood collection and biochemical analysis

The blood was collected via retro-orbital puncture from the overnight fasted mice prior to euthanasia. It was then allowed to clot at room temperature and centrifuged at 1500 g for 15 minutes at 4^0^C. The serum was collected and stored at −80^0^C until further analysis. Serum AST, ALT, cholesterol, triglycerides, and LDL-cholesterol were measured using the kits from Monlab tests, Spain, according to the manufacturer’s instructions.

### 2.9. Glucose tolerance test (GTT) and Insulin tolerance test (ITT)

The glucose tolerance test was performed on overnight fasted mice by measuring the baseline blood glucose level with the accu-check glucometer (Roche Diagnostics). Glucose (1g/kg body weight) dissolved in phosphate buffered saline was intraperitoneally injected into mice. After a glucose injection, blood glucose levels were measured at various time intervals (0, 15, 30, 60, 90, and 120 minutes). The insulin tolerance test was performed by fasting the mice for 5–6 h and measuring the baseline glucose levels similar to GTT. Mice were intraperitoneally injected with insulin (0.75 U/kg body weight). Blood glucose was measured thereafter at different time intervals (0, 15, 30, 45, 60, 90, and 120 minutes). GTT and ITT were performed before starting the compound treatment (12 weeks) and also at the end of the treatment (24 weeks).

### 2.10. Histological analysis

Paraffin-embedded formalin-fixed liver tissues were subjected to hematoxylin and eosin (H&E) staining to assess steatosis, hepatocyte ballooning and lobular inflammation. A pathologist at JSS Hospital scored the liver specimens (blindfolded) for severity of NAFLD (following the FLIP algorithm and NASH-Clinical Research Network (CRN) criteria) (Poynard et al., 2018; Brunt et al., 2019). Oil red o staining was performed on fresh liver sections to evaluate steatosis using established protocols.

### 2.11. TBARS assay

Lipid peroxidation was estimated by the TBARS assay following the manufacturer’s instructions. The production of TBARS was expressed as the equivalent of malondialdehyde (MDA). 25mg of liver tissue was weighed and homogenized. Tissue homogenate was used to measure the MDA levels. The reaction products were measured at 530nm excitation and 550nm emission. The MDA levels were normalized against standard values.

### 2.12. ROS assay

Intracellular reactive oxygen species production was determined in liver tissues by measuring 2’, 7’-dichlorofluorescein diacetate (DCFH-DA) (BioVision) following the manufacturer’s instructions. The single cell suspension was prepared from fresh liver tissue by enzyme digestion and the pelleted cells were incubated with ROX label for 45 minutes at 37°C in the dark. After incubation, fluorescence intensity was measured at excitation/emission= 495/529 nm using a multi-mode plate reader (EnSpire™ Multimode Plate Reader, Perkin Elmer) and the change in florescence was determined after background subtraction.

### 2.13. RNA isolation and quantitative real time PCR

Total RNA was isolated from frozen liver tissue using TRIzol (Sigma, USA). A Nano Drop spectrophotometer was used to determine the concentration and quality of RNA. The RNA was reverse transcribed into cDNA using the Verso cDNA synthesis kit (Thermo) following the manufacturer’s instructions. The Rotor-Gene Q 5plex HRM System (QIAGEN) was used to perform quantitative real-time PCR. The reactions were carried out using the DyNamo Colorflash SYBR green kit (Thermo), 0.5mM primers (IDT), and 50ng of cDNA in a 20µl reaction volume. The relative fold change in mRNA levels was calculated as 2^-ΔΔCt^ and was expressed normalized with endogenous control β-actin.

### 2.14. Western Blot

Liver tissue lysates were prepared by solubilizing the tissue in RIPA buffer (Sigma, USA) containing protease/phosphatase inhibitor (Thermo Fisher). Homogenized tissues were centrifuged at 13000 rpm for 10 minutes at 4^0^C and the supernatants were collected. The protein concentration of the tissue lysates was estimated using Bradford’s protein estimation method using Bio Rad protein Assay Dye Reagent (Bio Rad). To separate the proteins, 30 µg of protein samples were loaded into SDS-PAGE and transferred onto a nitrocellulose membrane for all western blots. The membranes were further blocked using 5% nonfat skim milk for 1 hour and probed with specific primary antibodies (pErk1/2, Erk1/2, pJNK, JNK, Santa Cruz) and β-actin (Cell Signaling). Upon overnight incubation at 4^0^C, membranes were incubated with secondary antibodies (Cell Signaling) for 2 hours at room temperature. The blots were visualized using the Western Bright ECL HRP substrate (Advansta). The blot images were analyzed using ImageJ software. The intensity of each band was normalized with endogenous control, β-actin.

### 2.15. Statistical Analysis

Results were calculated as means ± SEM. Statistical significance was analyzed using Student’s t-test for two groups or analysis of variance (ANOVA) with post hoc Bonferroni correction for multiple comparisons. All statistical analyses were performed using the GraphPad Prism software (version 6), and p values < 0.05 (*) or < 0.001 (**) were considered significant. In all cases, n=8-10 mice per group, unless otherwise indicated in the figure legends.

## 3. RESULTS

### 3.1. Quality evaluation and dose selection of *Tecomella undulata* for preclinical studies

As per the guidelines of ‘Quality control methods for medicinal plant materials’ by the World Health Organization (WHO) (https://www.who.int/publications/m/item/quality-control-methods-for-medicinalplant-materials) and the Ayurvedic Pharmacopoeia of India (API), Government of India (Joshi et al., 2017), the raw/crude stem bark powder of *Tecomella undulata* was subjected to pharmacopoeia tests. Various standardization components that were analyzed include macroscopic and microscopic characteristics, organoleptic characteristics, physiochemical properties, presence of residues (heavy metals, pesticides, toxins) and microbial load (**Supplementary Table 1**). After determining that the values were within the limits or matched with the pharmacopoeial standards of WHO and API, *Tecomella undulata* powder was used for the mice study. Furthermore, *in vitro* cell viability assays were also performed prior to preclinical studies.

The choice of *Tecomella undulata* dosage was determined based on the human therapeutic dose and body surface area conversion factor as described in the methods section. A dose of 80 mg/kg body weight of mice was chosen for the *in vivo* experiments.

### 3.2. *Tecomella undulata* treatment ameliorates western diet sugar water-induced obesity and insulin resistance

C57Bl/6 mice were divided into two groups and fed with chow diet normal water (CDNW) and western diet sugar water (WDSW), respectively. The mice were further divided into treatment groups for vehicle control, Saroglitazar (4 mg/kg body weight), and *Tecomella undulata* (80 mg/kg body weight). Mice fed with WDSW became obese with increased body weight and liver weight compared to mice fed with CDNW. Additionally, WDSW mice also developed insulin resistance. The mice treated with *Tecomella undulata* for 12 weeks showed a significant reduction in body weight and liver weight compared to vehicle control (**Figure 1A and 1B**). In addition, calorie intake was not affected by *Tecomella undulata* treatment, which indicates that the decrease of body weight upon treatment was not due to changes in food consumption. *Tecomella undulata* also significantly reduced fasting glucose and fasting insulin levels in WDSW fed mice (**Figure 1C and 1D**). Insulin resistance, a key driver of NASH was also reduced upon *Tecomella undulata* treatment as measured by HOMA-IR (homeostatic model Assessment for insulin resistance) (**Figure 1E**).

**Figure 1.**
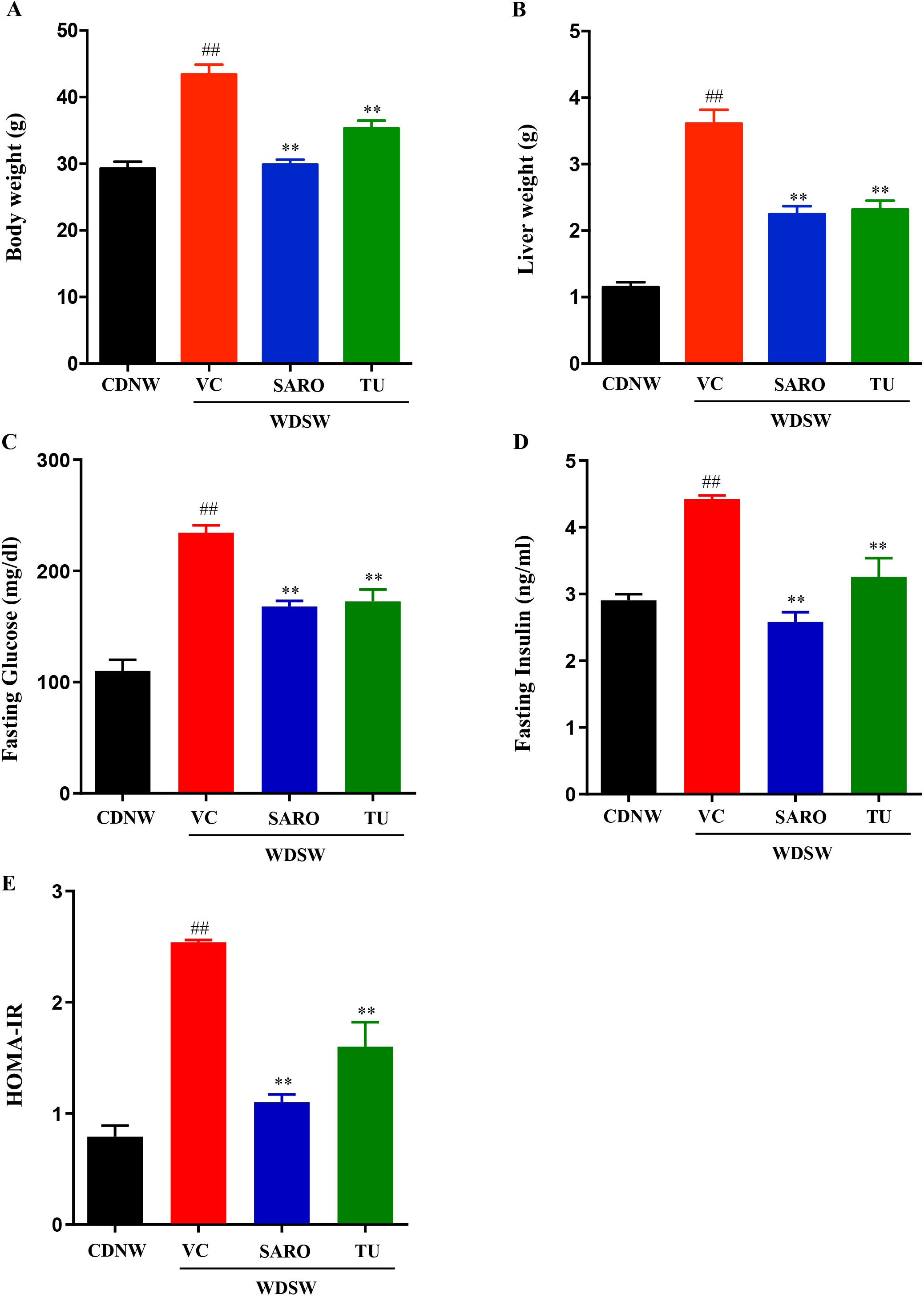
*Tecomella undulata* (TU) reduced body weight and improved insulin resistance in high fat diet-induced obese mice. Mice were treated with CDNW or WDSW for 12 weeks and following that; WDSW mice were divided into 3 groups and treated with (i) vehicle control (VC) (ii) Saroglitazar (SARO) (iii) *Tecomella undulata* (TU) via oral gavage for additional 12 weeks. At the completion of the treatment, body weights (**A**), liver weight (**B**), fasting glucose (**C**), and fasting insulin (**D**) were measured. HOMA-IR (**E**) was calculated using the formula: (fasting insulin (milliunits/liter)×fasting glucose (mmol/liter))/22.5. Data are expressed as mean±SEM for 8-10 mice per group. ##p < 0.001 compared to CDNW; **p < 0.001 compared to WDSW, vehicle control.

Additionally, administration of *Tecomella undulata* improved glucose tolerance (assessed by glucose tolerance test) and insulin sensitivity (assessed by insulin tolerance test) in hyperglycemic WDSW mice compared with vehicle alone (**Figure 2A and 2B**) but had no effect in CDNW mice. Overall, the degree of amelioration was on par with Saroglitazar, the positive control group used in the study.

**Figure 2.**
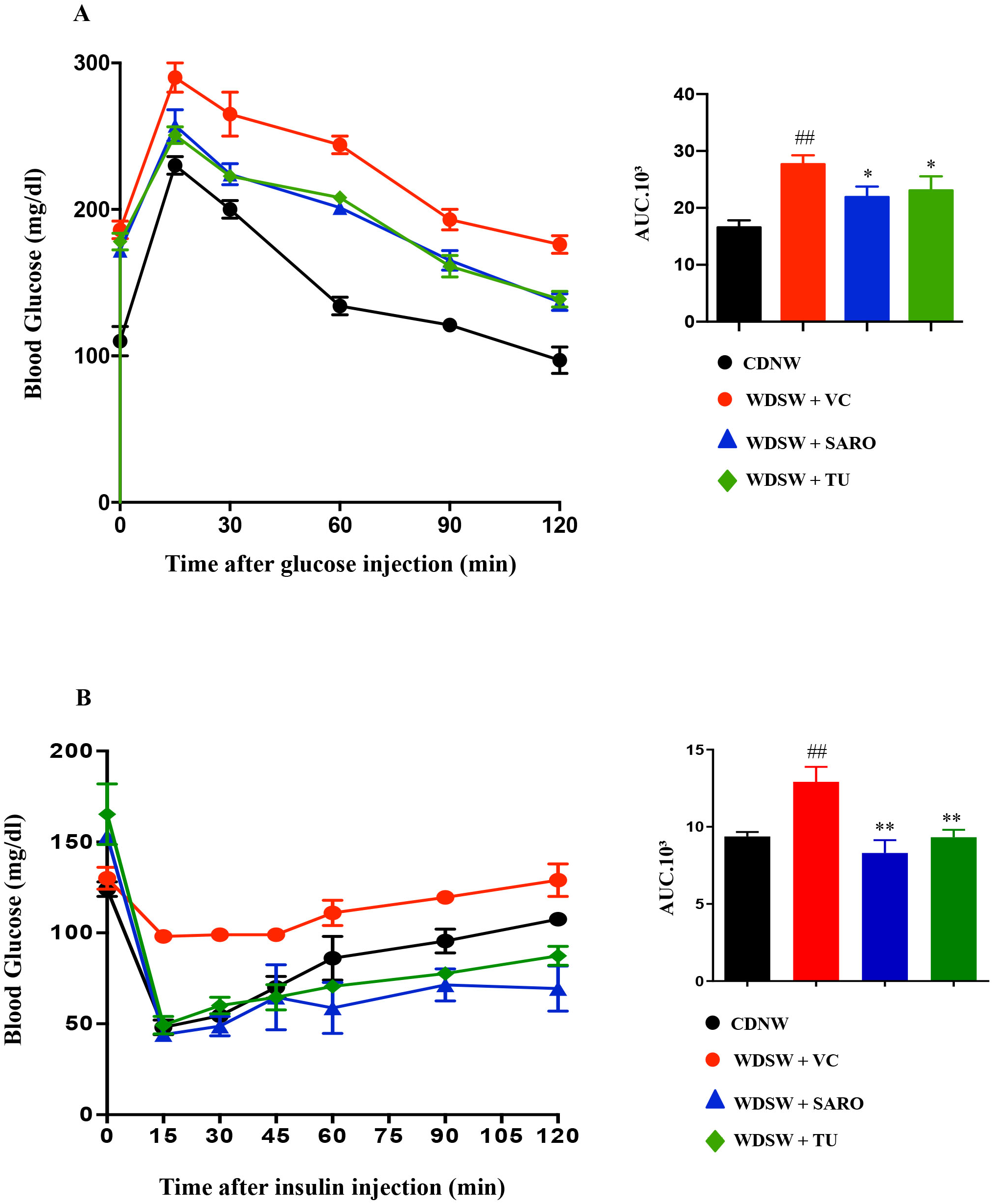
*Tecomella undulata* (TU) improves glucose tolerance and insulin sensitivity. CDNW and WDSW mice treated with vehicle control (VC), Saroglitazar (SARO) or *Tecomella undulata* (TU) for 12 weeks. Mice were fasted overnight and blood glucose concentration (mg/dl) was measured after intraperitoneal injection of 1g/kg glucose (**A**). Mice fasted for 4-5 hours were administered 0.75 units/kg insulin and blood glucose concentration (mg/dl) measured (**B**). The bar graphs depict the area under curve (AUC) with or without treatment. Data are expressed as mean±SEM for 8-10 mice per group. ##p < 0.001 compared to CDNW; **p < 0.001 or *p < 0.05 compared to WDSW, vehicle control.

### 3.3. *Tecomella undulata* prevented liver injury and hyperlipidemia

As expected, the WDSW diet drastically elevated liver enzymes and the mice developed hyperlipidemia, the major indicator of liver injury. However, *Tecomella undulata* treatment in these mice showed a reduction in the serum levels of ALT and AST (**Figure 3A and 3B**). In addition, *Tecomella undulata* administration also improved circulating triglycerides, total cholesterol, and LDL-cholesterol in WDSW mice compared to the vehicle control group (**Figure 3C-E**). Consistently, *Tecomella undulata* had no effect on the liver enzymes and lipid profile of CDNW mice as with Saroglitazar.

**Figure 3.**
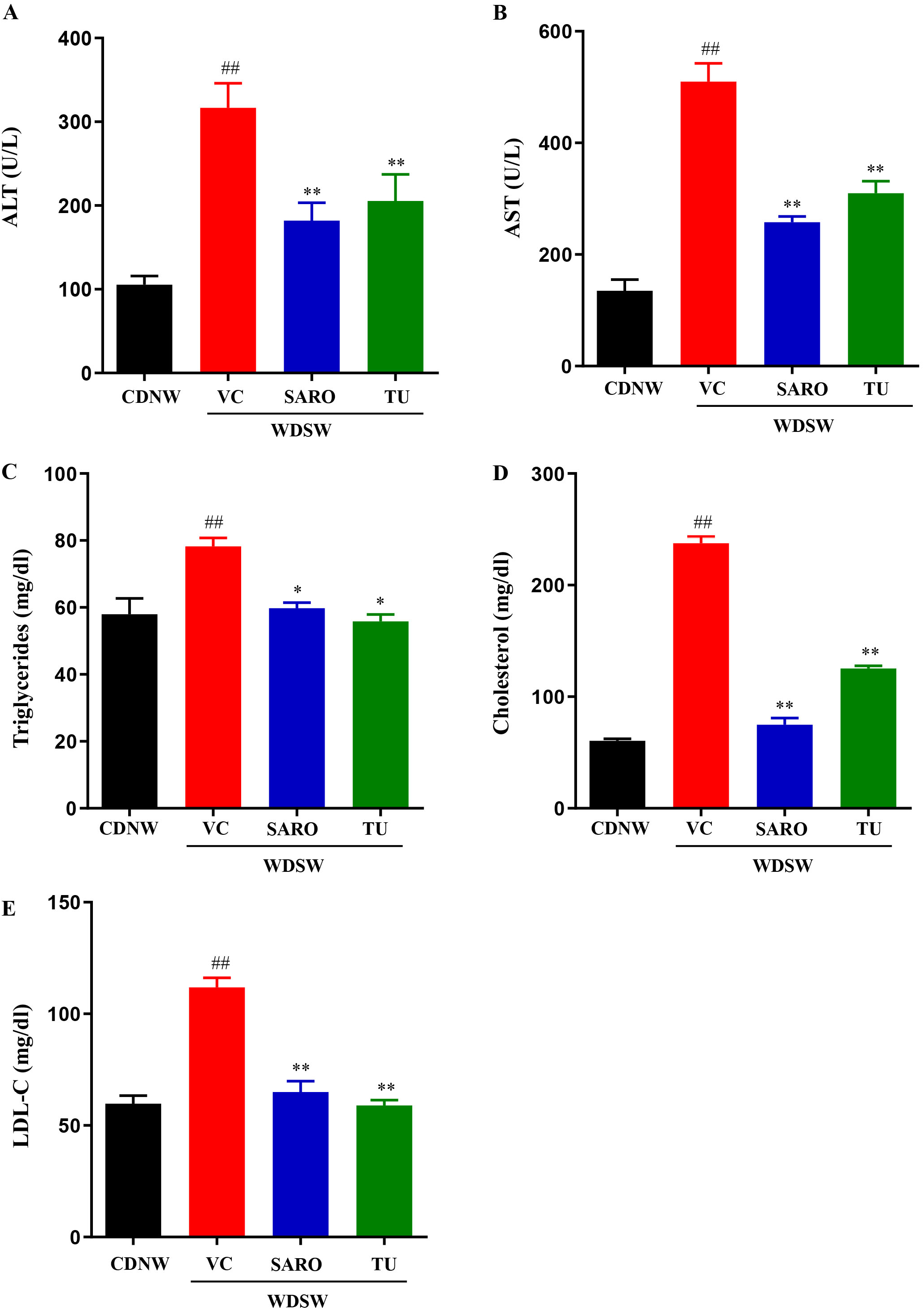
*Tecomella undulata* (TU) treatment reduced liver injury and hyperlipidemia. CDNW or WDSW for 12 weeks were administered vehicle control (VC), Saroglitazar (SARO) or *Tecomella undulata* (TU) for another 12 weeks. At the end of 24 weeks following dietary intervention and treatment, mice were fasted overnight and blood was collected. (A) serum ALT, (B) serum ALT, (C) serum triglycerides, (D) serum cholesterol (E) serum LDL-C. Data are expressed as the mean±SEM for 8–10 mice per group. ##p < 0.001 compared to CDNW; **p < 0.001 or *p < 0.05 compared to WDSW, vehicle control. AST, aspartate aminotransferase; ALT, alanine aminotransferase; LDL-C, low-density lipoprotein-Cholesterol.

Collectively, these data demonstrate that *Tecomella undulata* significantly ameliorates western diet sugar water-induced obesity, insulin resistance, liver injury and dyslipidemia.

### 3.4. *Tecomella undulata* improves hepatic steatosis and steatohepatitis

Histological analysis was carried out for all the groups. After 24 weeks, CDNW mice did not develop fatty liver, while WDSW mice with vehicle control developed grade 3 macrovesicular steatosis. The increased lipid droplets (both in number and size) were very much evident as evaluated by Hematoxylin and Eosin (H&E) staining in WDSW mice (**Figure 4A**), and *Tecomella undulata* treatment significantly reduced the number and size of the lipid droplets. Similar results were also confirmed by Oil Red O staining (**Figure 4B**). These WDSW mice showed significant steatosis (3±0.1), hepatocellular ballooning (2±0.2), and lobular inflammation (2.6±0.25) with the mean ± SEM NAFLD activity score (NAS) of 7.6±0.3 confirming the development of NASH (**Figure 5**).

**Figure 4.**
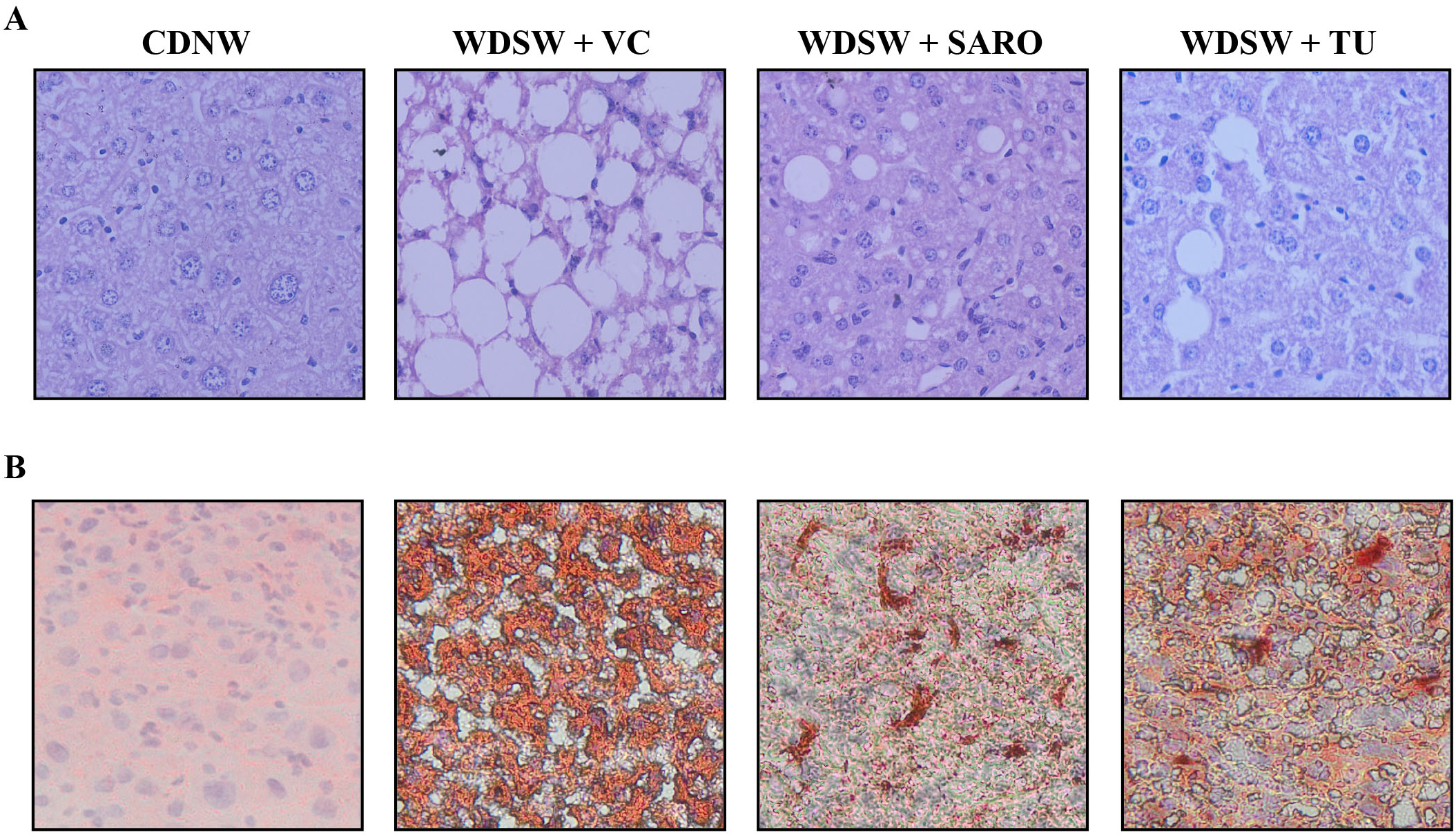
Histological features of mice treated with *Tecomella undulata* (TU). Representative microscopic views of liver sections from CDNW or WDSW mice treated with vehicle control (VC), Saroglitazar (SARO) or *Tecomella undulata* (TU). Intracellular lipid accumulation were determined by staining with (**A**) Hematoxylin and Eosin (H&E), and (**B**) Oil Red O.

**Figure 5.**
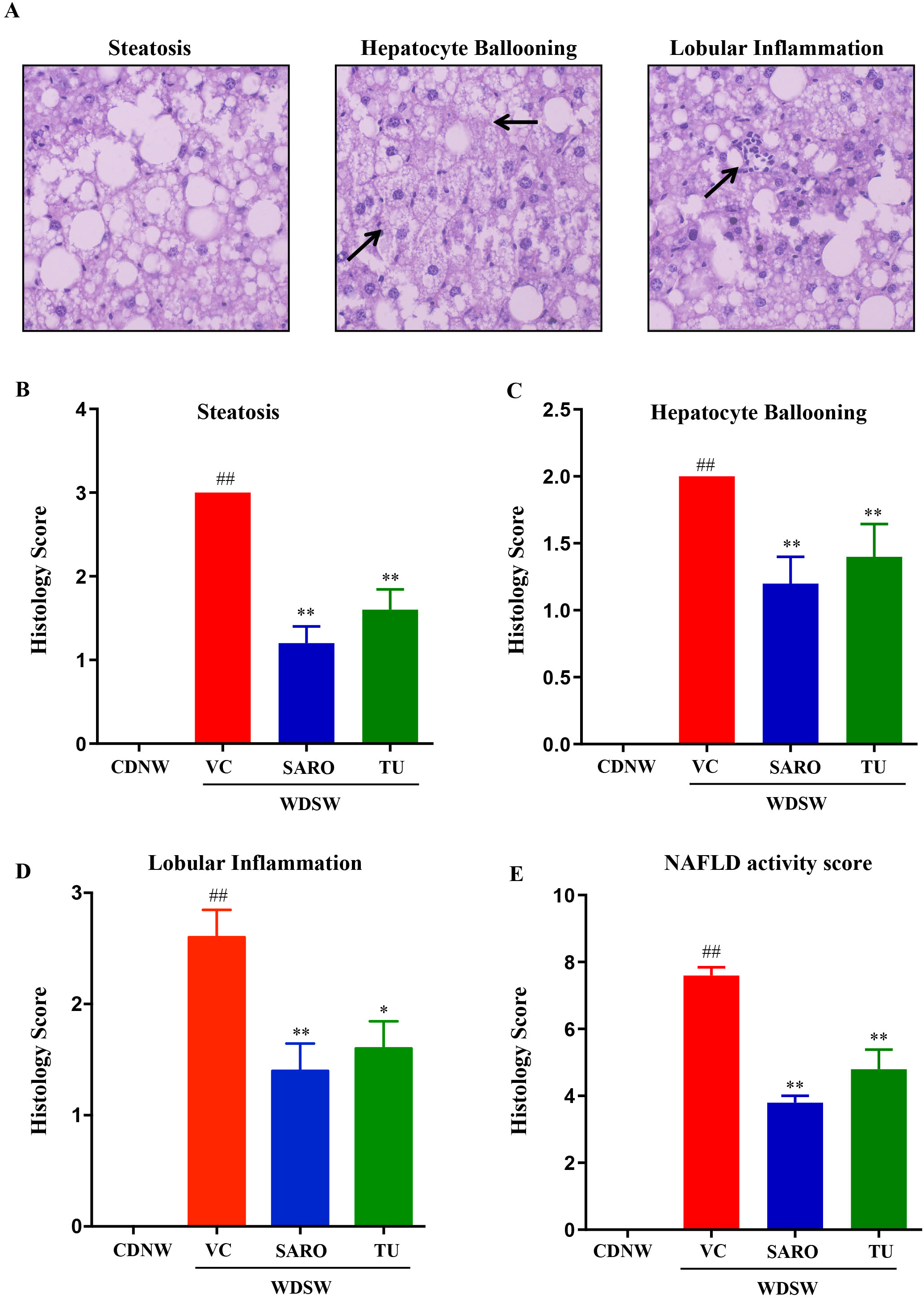
*Tecomella undulata* (TU) treatment ameliorates fatty liver and steatohepatitis. CDNW or WDSW mice treated with vehicle control (VC), Saroglitazar (SARO) or *Tecomella undulata* (TU) for 12 weeks. (**A**) Hematoxylin and Eosin (H&E) stained liver sections depicting steatosis, hepatocyte ballooning, and lobular inflammation. Histology score for (**B**) steatosis, (**C**) hepatocyte ballooning, (**D**) lobular inflammation, (**E**) NAFLD activity score were quantified. Data are expressed as the mean±SEM for 8–10 mice per group. ##p < 0.001 compared to CDNW; **p < 0.001 or *p < 0.05 compared to WDSW, vehicle control.

Of note, *Tecomella undulata* treatment significantly reduced steatosis (1.6±0.2), hepatocellular ballooning (1.4±0.25), and lobular inflammation (1.8±0.37). The NAFLD activity score in the *Tecomella undulata* treated mice was significantly lower (4.8±0.6) compared to the WDSW group treated with vehicle (**Figure 5B-5E**). All the histological scoring data were promising and were comparable to those of Saroglitazar, the positive control group.

### 3.5. *Tecomella undulata* alleviates hepatic ER stress and Oxidative stress

The pathogenesis of NASH is characterized by chronic impairment of lipid metabolism, which results in cellular lipotoxicity, endoplasmic reticulum (ER) stress, lipid peroxidation, and mitochondrial dysfunction (Parthasarathy et al., 2020). We examined whether administration of *Tecomella undulata* via oral gavage could prevent the progression of oxidative stress and ER stress, leading to improved liver function and reduced inflammation. WDSW mice showed a significant increase in ER stress as measured by the expression of markers such as C/EBP homologous protein (CHOP) and 78-kDa glucose regulated protein (Grp78). However, *Tecomella undulata* significantly reduced the expression level of ER stress markers (**Figure 6A and 6B**). Furthermore, oxidative stress was determined by measuring the levels of reactive oxygen species (ROS) and malondialdehyde (MDA, a product of lipid peroxidation) in the liver tissues of the experimental mice. As expected, *Tecomella undulata* treated mice showed reduced levels of ROS and MDA in comparison to the WDSW mice treated with vehicle control (**Figure 6C and 6D**). Consistently, we observed increased levels of cellular antioxidants as reflected by superoxide dismutase (SOD) and catalase (CAT) expression (**Figure 6E and 6F**) in *Tecomella undulata* treated WDSW mice compared to vehicle control.

**Figure 6.**
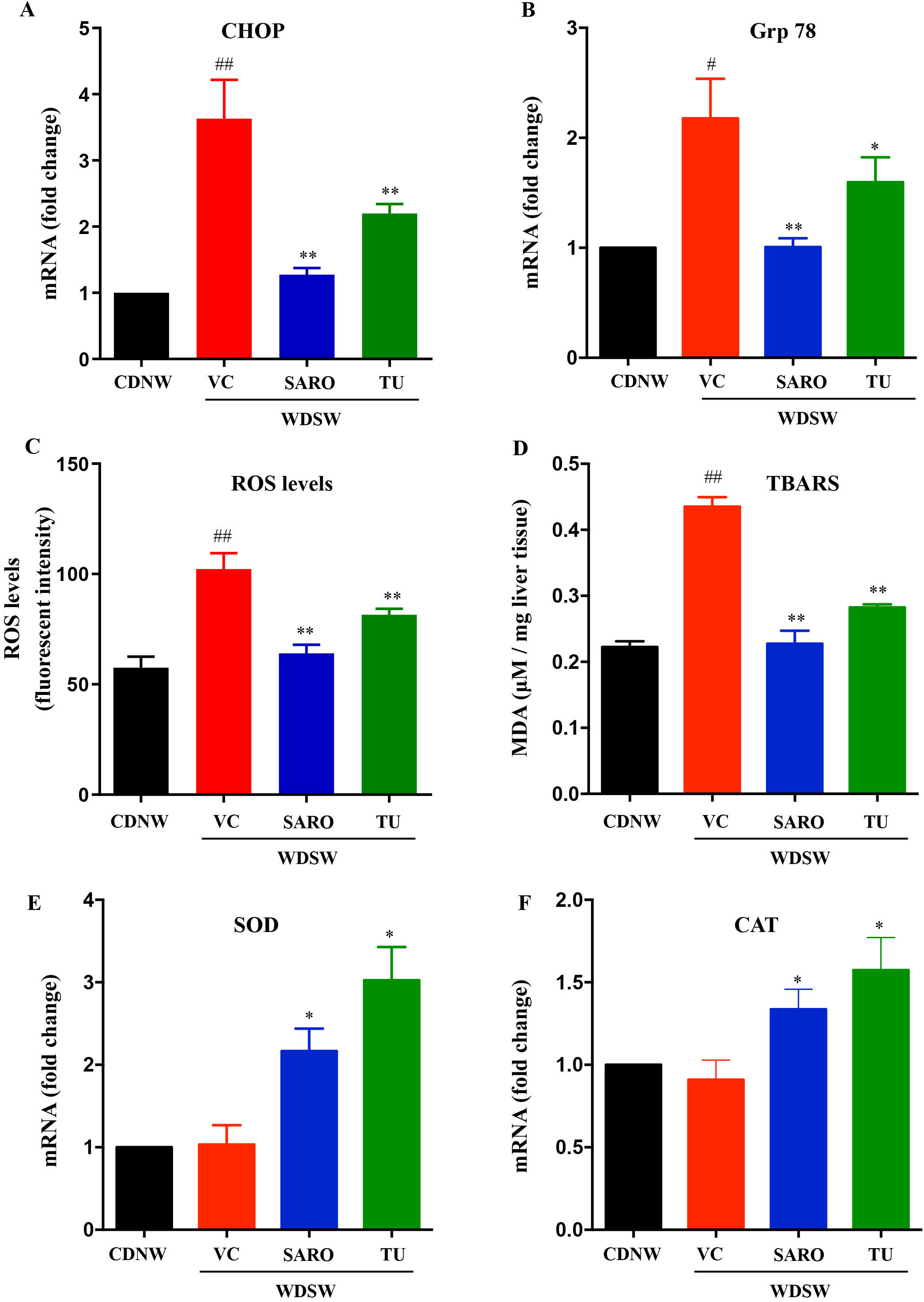
Effect of *Tecomella undulata* (TU) on ER stress and oxidative stress in NASH mice. The relative expression of hepatic mRNA levels of CHOP (**A**), Grp78 (**B**), SOD (**E**), and CAT (**F**) were determined using qRT-PCR. The experiments were carried out in triplicates and β-actin was used as endogenous control for normalizing the mRNA levels. Intracellular ROS production (**C**) and lipid peroxidation (MDA levels) (**D**) was determined in the liver tissues of CDNW and WDSW mice treated with vehicle control (VC), Saroglitazar (SARO) or *Tecomella undulata* (TU). Data are expressed as the mean±SEM for 8–10 mice per group. ##p < 0.001 or #p < 0.05 compared to CDNW; **p < 0.001 or *p < 0.05 compared to WDSW, vehicle control. CHOP, C/EBP homologous protein; Grp78, 78 kDa glucose regulated protein; SOD, superoxide dismutase; CAT, catalase; ROS, reactive oxygen species; MDA, malondialdehyde.

### 3.6. Anti-inflammatory effects of *Tecomella undulata* in NASH

Since inflammation plays a key role in the pathogenesis and progression of NASH, modulation of the antioxidant responses may serve as a target to prevent the disease (Tilg and Moschen, 2010). In this study, we investigated the proinflammatory markers, tumor necrosis factor α (TNFα) and interleukin-1β (IL-1β) and found that *Tecomella undulata* administration to WDSW mice reduced inflammation (**Figure 7A and 7B**). Consistently, the activated and phosphorylated c-Jun N-terminal kinase (JNK) and extracellular signal-regulated protein kinase (ERK1/2), which are primary stress kinases that play a fundamental role in the development of steatosis and are important players in inducing inflammation (Urano et al., 2000; Khan et al., 2017), were also found to be downregulated upon *Tecomella undulata* treatment (**Figure 7C and 7D**). All this data was comparable to Saroglitazar.

**Figure 7.**
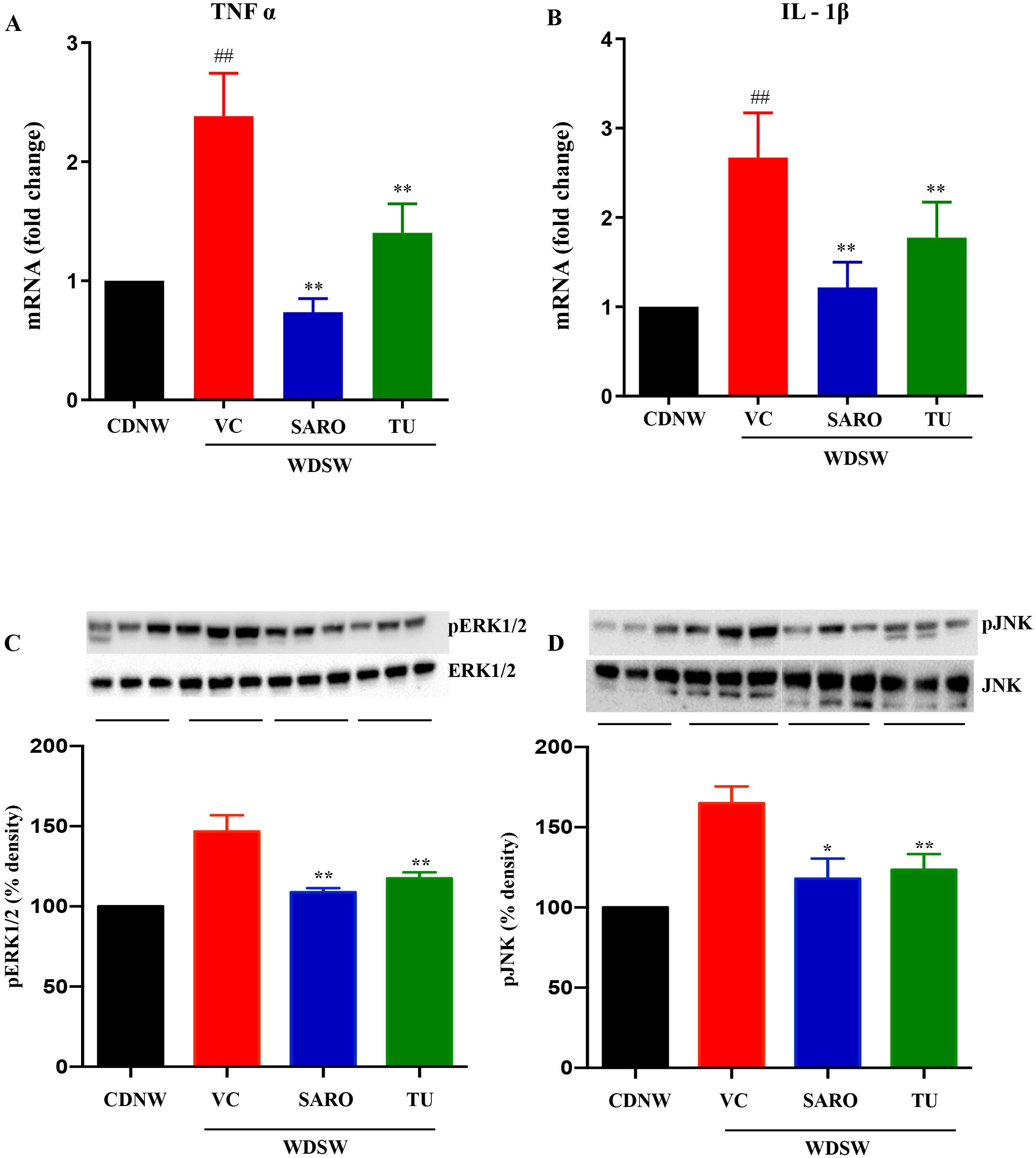
*Tecomella undulata* (TU) treatment ameliorated hepatic inflammation in NASH mice. The relative expression of hepatic mRNA levels of TNFα (**A**), and IL-1β (**B**) were determined using qRT-PCR. The experiments were carried out in triplicates and β-actin was used as endogenous control for normalizing the mRNA levels. Whole cell lysates were prepared from liver tissue of CDNW and WDSW treated with vehicle control (VC), Saroglitazar (SARO) or *Tecomella undulata* (TU) for 12 weeks. Immunoblot analyses were performed for p-Erk1/2 and Erk1/2 (**C**), and p-JNK and JNK (**D**). Bar graphs show the densitometric values calculated after normalization to total Erk1/2 or total JNK. Data are expressed as the mean±SEM. ##p < 0.001 compared to CDNW; **p < 0.001 compared to WDSW, vehicle control. TNFα, tumor necrosis factor α; IL-1β, interleukin-1β; p-Erk1/2, phospho-extracellular signal-regulated protein kinase; p-JNK, phosphor-c-Jun N-terminal kinase.

**Figure 8.**
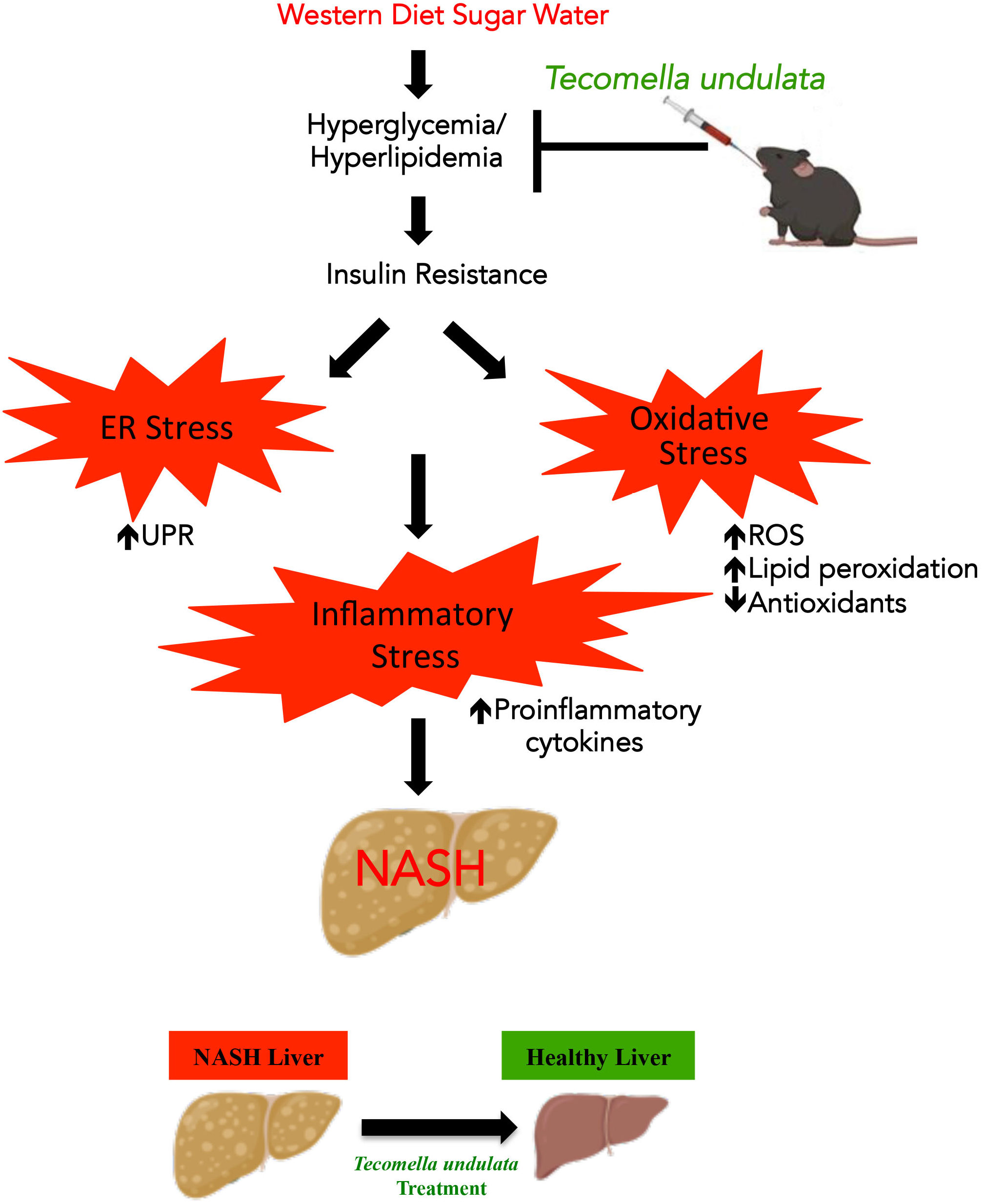
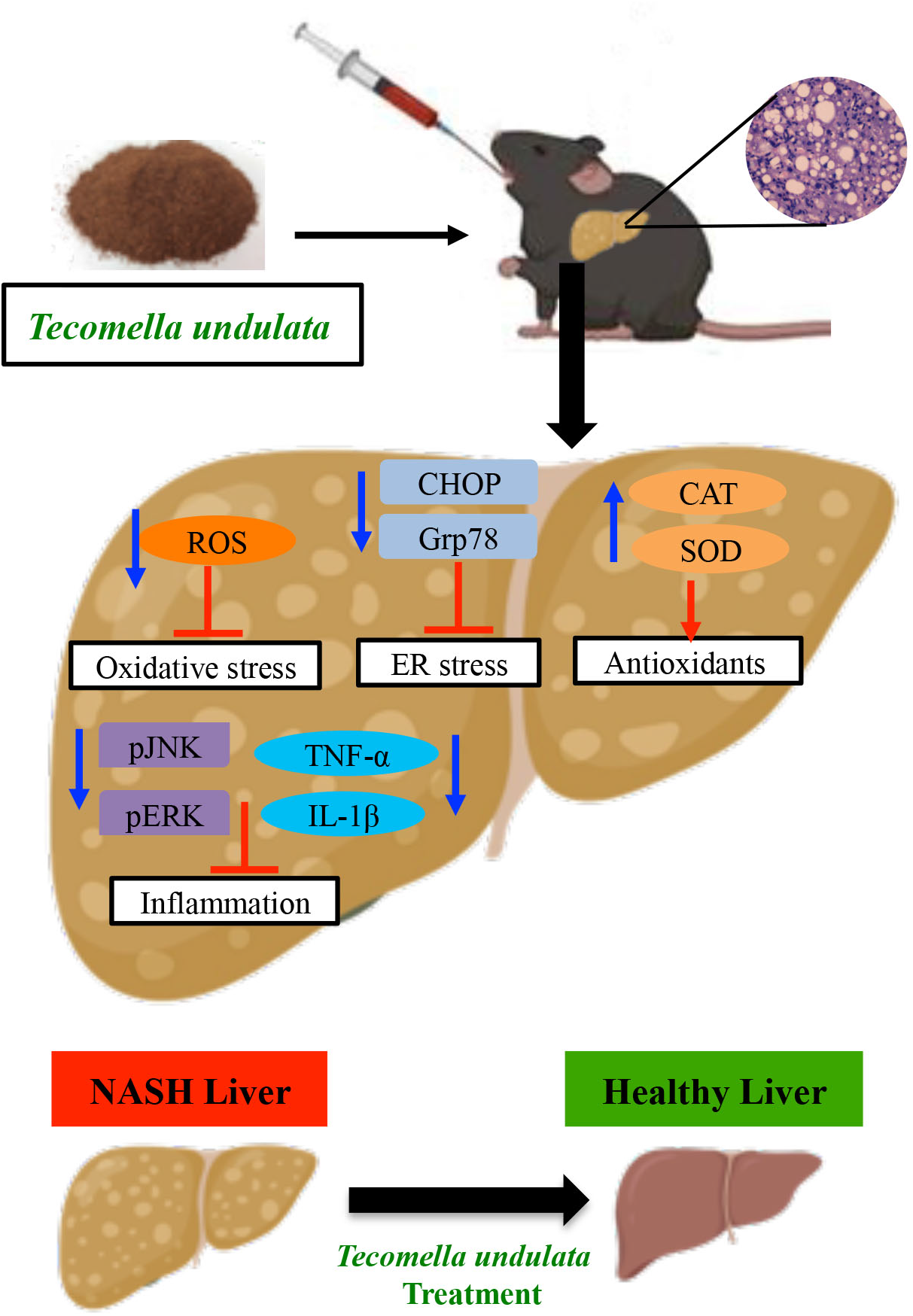
Schematic representation depicting the mechanisms involved in *Tecomella undulata* (TU) -mediated amelioration of NASH. Mice fed on WDSW for 12 weeks were treated with *Tecomella undulata* via oral gavage for another 12 weeks. Administration of *Tecomella undulata* ameliorated steatohepatitis in WDSW mice by improving insulin sensitivity, ER stress, oxidative stress, and increased production of antioxidants providing evidence that *Tecomella undulata* may have the potential to be used in the treatment of NASH.

Taken together, our results showed that *Tecomella undulata* could alleviate the western diet sugar water-induced ER stress and oxidative stress, improve liver injury and antioxidant status, and thereby reduce inflammation.

## 4. DISCUSSION

Steatohepatitis, a progressive form of NAFLD, is characterized by severe steatosis, inflammation, and cellular stress. The spectrum further continues to different stages of fibrosis and end-stage consequences like cirrhosis or hepatocellular carcinoma (Hardy et al., 2016). The primary treatment regimen focuses on NASH resolution, which includes several drug candidates, as well as regular physical activity and caloric restriction (Younossi et al., 2018). Due to NASH’s complexity and multifaceted pathological nature, there is an unmet need for the continued identification of novel therapeutics (Neuschwander-Tetri, 2020). Hence, the use of herbal medications may therefore be a promising alternative with the potential for effective therapeutic benefit. To date, a number of plant extracts, polyherbal formulations, and phytochemicals have been examined for their potential therapeutic value in the treatment of various liver diseases, including NASH (Jadeja et al., 2014). The current study provides proof of concept that *Tecomella undulata* improves histological characteristics as well as the key biological processes that are associated with the pathogenesis of NASH.

Despite the fact that there is literature on the preclinical evaluation of herbal remedies for the treatment of NASH, the primary flaw in the study lies in not making the right choice of the animal model. The quality of the diet, animal strain, concentration of fat, cholesterol, and sugar, time period, and reproducibility are the key characteristics that are always overlooked in NAFLD research. There are a wide variety of genetic and dietary animal models that are available to study NASH (Santhekadur et al., 2018). However, choosing the ideal animal model mimicking human NASH with respect to diet-induced obesity and insulin resistance is crucial without much alteration in the macronutrient composition (deficiency or overnutrition) or the addition of toxins. In this study, we have used a mouse model (detailed in the methods section) that not only recapitulates the effects of the diet but also demonstrates concordance with histology, disease progression, biological processes, and pathways. With the exception of the fact that steatohepatitis takes around 24 weeks to manifest, the mice exhibit significant translatability to human NASH as they progressively develop obesity, insulin resistance, dyslipidemia, and inflammation. In addition to the experimental NASH model, other crucial elements, including dosage, timing of the intervention, route of administration, duration of exposure, and endpoint assessments, were thoroughly specified to assess the efficacy of *Tecomella undulata* for the treatment of NASH.

In this study, we provide *in vivo* evidence for the hepatoprotective effects of *Tecomella undulata* in ameliorating NASH. *Tecomella undulata* is known to have a special place in herbal medicine and has been widely recommended to treat hepatosplenomegaly, hepatitis, and obesity (Jain et al., 2012; Alvala et al., 2013). Studies have investigated the pharmacological effects of *Tecomella undulata* and have reported their hepatoprotective, anti-inflammatory, immunomodulatory, and antimicrobial activities (Goyal et al., 2012; Ahmed et al., 1994; Choudhary, 2011). However, there is a lack of scientific data illuminating the molecular processes through which *Tecomella undulata* exerts its effects in preclinical research using the appropriate experimental model. To our knowledge, this is the first study to investigate the effect of *Tecomella undulata* on hepatic damage caused by western diet sugar water. We evaluated the protective effects of *Tecomella undulata* against hepatic lipid accumulation in an experimental NASH model and how it regulates the genes and proteins involved in biological processes associated with the development and progression of NASH. The C57bl/6 mice develop fatty liver after 12 weeks following the initiation of a western diet sugar water regimen and progress to NASH in concordant with human NASH. The administration of *Tecomella undulata* to the mice showed improvement in obesity, as marked by a reduction in body weight and liver weight. There was also a significant amelioration in insulin resistance. Furthermore, we also demonstrated a marked improvement in the histological and biochemical outcomes upon treatment with *Tecomella undulata* in NASH mice. Collectively, it was interesting to note that the hepatoprotective effects of *Tecomella undulata* followed a similar trend to Saroglitazar, a dual peroxisome proliferator activated receptor α/γ agonist approved for the treatment of human NASH (Goyal et al., 2020, Kumar DP et al., 2020).

Previous studies have shown the hepatoprotective activity of ethanolic and methanolic extracts of *Tecomella undulata* in rats against thioacetamide- and carbon tetrachloride-induced hepatocyte damage, respectively (Khatri et al., 2009; Jain et al., 2012). Ethanolic extracts of *Tecomella undulata* have shown anti-hyperglycemic and antioxidant potential in streptozotocin-nicotinamide-induced type 2 diabetic rats, although the mechanism remains ill defined (Kumar et al., 2012). *In vitro* and *in vivo* studies of *Tecomella undulata* ethyl acetate extract revealed anti-obesity efficacy by downregulating adipogenesis and lipogenesis via sirtuin 1 (SIRT1), adiponectin, and peroxisome proliferator activated receptor γ (PPARγ) (Alvala et al., 2013). However, the current study provides insight into the mechanisms by which *Tecomella undulata* may mitigate NASH. Hepatocellular stress plays a prominent role in the onset of NASH. The large influx of free fatty acids saturates the oxidation mechanisms, leading to the production of reactive oxygen species (ROS), which are not neutralized by antioxidants, resulting in hepatic injury via the Ras/Raf/Erk pathway (Khan et al., 2017; Kamata et al., 2005). Toxic lipid species, such as palmitic acid, accumulate in the liver and cause ER stress, as evidenced by the unfolded protein response (UPR) (Lake et al., 2014; Song et al., 2019). Thus, the markers for ER stress (Grp78 and CHOP) and oxidative stress (ROS and MDA levels) were studied along with ERK1/2 and we found that treatment with *Tecomella undulata* reduced oxidative stress and ER stress in NASH mice, providing a rationale for low cholesterol and lipid reduction in the liver. As expected, the antioxidant levels were increased due to the treatment, as shown by SOD1 and CAT expression. The hepatic inflammatory response is a major driver of NAFL progression to NASH and fibrosis. Several intra- and extrahepatic factors trigger the inflammatory system, which exacerbates cellular injury and apoptotic response. Along with inflammatory mediators such as TNF-α and IL-1β, free cholesterol and oxidative stress can activate JNK-dependent proinflammatory pathways (Duan et al., 2022; Osto et al., 2008; Malhi et al., 2006). Our study reports the anti-inflammatory potential of *Tecomella undulata* by the marked reduction in cellular mediators of inflammation.

This study adds to the growing literature on the use of herbal medicines and supports the further assessment of *Tecomella undulata* in humans for the treatment of NASH. Previous studies have used analytical HPLC, which has led to the identification of betulinic acid, ferulic acid, rutin, quercetin, vanillic acid, and other nine chemicals in methanol or ethanol extract of *Tecomella undulata’s* stem bark (Jain et al., 2012, Alvala et al., 2013, Ali et al., 2017) but unfortunately no validation has been done in NASH. The present study sets the right stage in providing the strongest evidence for the traditional use of *Tecomella undulata* in herbal medicine with well-designed reverse pharmacology-guided research proving the safety and therapeutic efficacy. Based on the previous HPLC studies, future experiments are in the pipeline to isolate and purify potent hit compounds and also perform therapeutic validation of these pure compounds in the treatment of NASH rather limiting to just extract from the stem bark of *Tecomella undulata*. We also plan to conduct large scale randomized controlled trials (RCT) to support the therapeutic effects of Tecomella undulata in NASH. We strongly believe that the application of omics analysis (transcriptomics, proteomics, lipidomics, metabolomics, metagenomics) may also aid in exploring the molecular mechanisms in our studies.

## 5. CONCLUSION

In summary, we first discovered the promising hepatoprotective effects of the stem bark powder of *Tecomella undulata* in a preclinical model of NASH that closely mimics the dietary pattern, systemic milieu, and histological spectrum of NASH and also shows activation of the key cellular pathways linked to human disease. Our study demonstrated that *Tecomella undulata* improves insulin sensitivity by attenuating oxidative stress, ER stress, and inflammation in steatohepatitis. Thus, *Tecomella* undulata have a therapeutic role in the prevention of NASH.

## ABBREVIATIONS

NAFLD: nonalcoholic fatty liver disease
NASH: nonalcoholic steatohepatitis
ROS: reactive oxygen species
ER: endoplasmic reticulum
CD: chow diet
WD: western diet
NW: normal water
SW: sugar water
CMC: carboxymethylcellulose
AST: aspartate transaminase
ALT: alanine transaminase
LDL: low density lipoprotein
GTT: glucose tolerance test
ITT: insulin tolerance test
TBARS: thiobarbituric acid reactive substances
MDA: malondialdehyde
HOMA-IR: homeostatic model assessment for insulin resistance
NAS: NAFLD activity score
CHOP: C/EBP homologous protein
Grp78: 78-kDa glucose regulated protein
SOD: superoxide dismutase
CAT: catalase
TNFα: tumor necrosis factor α
IL-1β: interleukin-1β
JNK: c-Jun N-terminal kinase
ERK1/2: extracellular signal-regulated protein kinase
SIRT1: sirtuin 1
UPR: unfolded protein response
PPAR γ: peroxisome proliferator activated receptor γ
VC: vehicle control
SARO: Saroglitazar
TU: Tecomella undulata
SEM: standard error mean.

## ACKNOWLEDGMENT

We thank Dr. Puneshwar Keshari, Department of Ayurveda and Alternative Medicine, Kathmandu, Nepal for the valuable discussion and experiments on using *Tecomella undulata* for *in vivo* study.

## AUTHOR CONTRIBUTIONS

DeS, PP, S, and DPK conceptualized the project; ANS, SG, SBC, and DPK designed the study; ANS, DS, and DPK performed the experiments and analyzed the data; ANS, DeS, PP, S, SS, SG, and DPK interpreted the data; and ANS and DPK wrote the manuscript. All authors contributed to the article and approved the submitted manuscript.

## FIGURE AND TABLE LEGENDS

**Supplementary Table 1. Results of Pharmacopeia analysis on *Tecomella undulata* (TU) stem bark powder sample.**

